# Environmental similarity between relatives reduces heritability of reproductive timing in wild great tits

**DOI:** 10.1101/2024.03.11.584392

**Authors:** Carys V. Jones, Charlotte E. Regan, Josh A. Firth, Eleanor F. Cole, Ben C. Sheldon

## Abstract

Intraspecific variation is necessary for evolutionary change and population resilience, but the extent to which it contributes to either depends on the causes of this variation. Understanding the causes of individual variation in traits involved with reproductive timing is important in the face of environmental change, especially in systems where reproduction must coincide with seasonal resource availability. However, separating the genetic and environmental causes of variation is not straightforward, and there has been limited consideration of how small-scale environmental effects might lead to similarities between individuals that occupy similar environments, potentially biasing estimates of genetic heritability. In ecological systems, environments can often be complex in their spatial structure, and it may therefore be important to account for similarities in the environments experienced by individuals within a population beyond considering spatial distances alone. Here, we construct multi-matrix quantitative genetic animal models using over 11,000 breeding records (spanning 35 generations) of individually-marked great tits (*Parus major*) and information about breeding proximity and habitat characteristics to quantify the drivers of variability in two key seasonal reproductive timing traits. We show that the environment experienced by related individuals explains around a fifth of the variation seen in reproductive timing, and accounting for this leads to decreased estimates of heritability. Our results thus demonstrate that environmental sharing between relatives can strongly affect estimates of heritability and therefore alter our expectations of the evolutionary response to selection.

## Introduction

Changing environments, resulting from a combination of biotic and abiotic processes, but particularly accelerated by human influences (IPCC, 2022) pose a challenge for individuals and populations in today’s world. Most of the traits that determine success in the face of such challenges are likely to be continuously distributed quantitative traits. To understand the evolutionary causes and consequences of intraspecific variation in quantitative traits it is necessary to estimate additive genetic variance and heritability (Falconer and Mackay, 1996). Narrow-sense heritability describes the proportional contribution of additive genetic variance to the observed phenotypic variance; its mis-estimation can lead to erroneous conclusions about a population’s evolutionary potential and resilience. Quantitative genetic methods, developed initially for animal and plant breeding are now applied widely to wild populations, using the resemblance of phenotypes between relatives, along with the consideration of environment effects that contribute towards variation, to estimate the heritability of a trait (Bonnet et al., 2022; Falconer and Mackay, 1996; Kruuk, 2004; Lynch and Walsh, 1998; Postma and Charmantier, 2007; Wilson et al., 2010).

Understanding the causes of variation in traits that are associated with timing (i.e. ‘phenological traits’) is particularly important as they are closely associated with the environment (Forrest and Miller-Rushing, 2010; Pau et al., 2011), and often show considerable variation across time and space, which is maintained despite close links with reproductive success (Reed et al., 2010). Phenology encompasses a wide range of seasonal timing traits, from breeding timing in birds, to hibernation in mammals, and flowering in plants; most phenological traits show continuous variation across individuals within the population (Cole and Sheldon, 2017; Germain et al., 2016; Matthysen et al., 2021). Understanding the causes of this individual variation, and accurately estimating the heritability of phenological traits is increasingly important with accelerating global change (Forrest and Miller-Rushing, 2010). For organisms that breed seasonally, selection is often expected to favour timing events to coincide with temporally varying resources in other trophic levels (Kharouba and Wolkovich, 2020; Park and Post, 2022; Perrins, 1969; Renner and Zohner, 2018; Samplonius et al., 2021). As such, timing has likely consequences for breeding success and survival and is therefore an important life history trait to understand in the context of environmental change (Brook et al., 2015; Simmonds et al., 2020; Thomas et al., 2001).

Variation in phenological traits have been linked to various abiotic factors at a range of temporal and spatial scales including altitudinal and latitudinal gradients, climatic conditions, habitat quality and food availability (Lane et al., 2018; Réale et al., 2003; Rubolini et al., 2007; Wilkin et al., 2007). In particular, increasing temperatures have been strongly linked to advancement in breeding time for a number of species (Both et al., 2004; Fitter et al., 1995; Moyes et al., 2011; Tryjanowski et al., 2003). Most frequently, these phenological shifts have been demonstrated at the population level by analysing changes in mean phenotypes in relation to a mean measure of the environment. However, for many organisms, the assumption that all individuals within a population experience the same environment is an oversimplification and will likely lead to inaccurate estimation of the relative importance of additive genetic and environmental effects to phenological variation. Therefore, quantifying individual level patterns in phenology is vital for understanding the population level patterns (Cole et al., 2021; Gervais et al., 2022), and the capacity for adaptation to climate change (Houle 1992; Forrest and Miller-Rushing 2010; Charmantier et al. 2014).

In natural populations the environment may be highly heterogenous on many dimensions, so individuals within populations will experience different environmental conditions, contributing to the observed variation in phenotypes. Individuals that share an environment may have more similar phenotypes, known as ‘common environment’ effects. Furthermore, relatives (i.e. individuals who share genes) may also often share environments due to factors such as limited dispersal, inheritance of breeding locations, habitat imprinting, maternal effects and temporal overlap (Davis and Stamps, 2004; Van Der Jeugd and McCleery, 2002). Animal model approaches have historically considered only a few key sources of common environment effects, most commonly maternal identity, birth year or habitat type (e.g. (Liedvogel et al., 2012; McCleery et al., 2004; Wilson et al., 2005). Not accounting for these shared environments among individuals risks assuming that observed phenotypic similarity is due to shared genes, and previous research has suggested this may lead to an upward bias in estimates of heritability (Gervais et al., 2022; Kruuk and Hadfield, 2007; Regan et al., 2017; Rutschmann et al., 2020; Stopher et al., 2012).

A small number of studies have looked at space sharing and its effects on heritability estimates (Gervais et al., 2022; Regan et al., 2017; Rutschmann et al., 2020; Stopher et al., 2012). Whilst these approaches give an improved picture, most previous studies assumed that spatial proximity is the most important factor, or that spatial proximity is a good proxy for environmental similarity. As such, little work has actually considered the true contribution of environmental similarity between individuals not close in space, or individuals that are close in space but subject to different environmental conditions. Indeed, while in many ecological systems the expectation may be that places closer together will be more similar, this baseline assumption may not always be correct, particularly given that environments are often complex in their spatial structure over distances relevant to the scales that individual organisms operate over. Therefore, to accurately estimate the additive genetic contribution to phenotypic variation there must be consideration of how phenotypes are influenced by similarities in the environment (both biotic and abiotic) experienced by individuals at an appropriate scale, regardless of their spatial proximity. For example, it is easy to conceive of arrangements of patchy environments such that individuals close together may experience very different environments, whilst those further apart may actually experience more similar environments. These sorts of common environment effects will be important for understanding how and when heritability estimates may be biased (Gervais et al., 2022; Thomson et al., 2018), but accounting for environmental similarity within quantitative genetic methods is challenging and thus has been less explored.

A popular model system for exploring genetic and environmental contributions to phenology has been breeding time in birds, normally studied as the date the first egg of the clutch is laid, or the date that the first egg(s) in the clutch hatches. Estimates of heritability of breeding time in birds range from 0.001 to 0.45 (Teplitsky et al., 2010; van Noordwijk et al., 1981), although relatively few studies have explicitly explored the role of shared environments, other than by fitting grouping variables to control for these. Compared to other reproductive traits in birds, like clutch size, breeding timing shows lower heritability, but greater variation between and within years, suggesting there is a larger influence of environmental factors (Evans et al., 2020; Van Der Jeugd and McCleery, 2002). We expect breeding timing to be closely tied to the environment, as individuals must rely on phenological cues to provide information on when there will be food available to feed their young.

In this study we assess the quantitative genetics of variation in breeding time in great tits studied at Wytham woods near Oxford over the past 62 years. Previous work has estimated heritability of timing in this population (Evans et al., 2020; Garant et al., 2008; Liedvogel et al., 2012; McCleery et al., 2004; Van Der Jeugd and McCleery, 2002), but has not addressed the spatial and environmental determinants of timing and the effect of their inclusion on heritability. Here we substantially extend this work by partitioning the variance in two traits associated with breeding timing (laying and hatching date) whilst accounting for different aspects of shared environments. Our aims were (1) to quantify how much between-individual variation is due to spatial and breeding environment factors, and how correlated breeding environmental similarity and spatial autocorrelation measures are, and (2) establish how including these factors in the models impacts heritability estimates.

## Results

### Initial findings

An influential previous analysis of these data used mother-daughter regression split by three dispersal classes to demonstrate environmental dependence of heritability (van der Jeugd and McCleery 2002). For completeness, we show that the findings in Van Der Jeugd and McCleery, (2002) are robust to repeated analysis using parent-offspring regressions with a substantially increased data set (see supplementary information Section 1). Since this approach does not make full use of the relatedness structure and doesn’t allow modelling of continuous environmental distance, we focus from here onwards on results using multi-matrix animal models.

Visually comparing the spatial proximity and environmental similarity matrices suggests that they are correlated at short distances but that this correlation declines as distance increases (Supplementary Figure 5). In line with this, mantel tests showed a correlation of 0.192 between the spatial proximity and breeding environment similarity matrices.

### Laying date

As expected, breeding year explained a considerable amount of variation in laying date in all models (ranging from 48.8 *±* 8.8% to 60.2 *±* 10.9%, see Table 1 and Figure 1A). The nest box model significantly improved the model fit compared to the minimal model without the nest box effect (χ2 =237, p < 0.001), with the nest box random effect explaining almost 3% of the variation (Figure 4a). Within-year heritability was very similar between these two models; adding the nest-box random effect decreased the estimate by 0.7 percentage points, corresponding to a change of 3.5% (minimal model: 20.1 *±* 5.8%, nestbox model: 19.4 *±* 5.4%: Table 1 and Figure 1C). Fixed effects are reported in Supplementary Table 5. The effect of mother’s age at breeding did not change much across models.

**Table 1.** Variance components from animal models (models 1-4). Details of models found in methods. Est (SE) gives the raw estimate and standard error. Rel to *V*P is the ratio of each variance component to *V*_P_. *V*_BY_ = breeding year, *V*_PE_ = focal individual permanent environment effect, *V*_A_ = focal individual additive genetic effect, *V*_NB_ = nest box random effect, *V*_SPATIAL_ = spatial proximity matrix_"_ *V*_BREED. ENV_. = breeding environment similarity matrix, *V*_R_ = residual variance, *V*_P_ = total phenotypic variance as sum of all variance component, *V*_P-within year_ = total phenotypic variance as sum of all variance component excluding *V*_BY,_ *h*^2^ = within year heritability as proportion of *V*_A_ to *V*_P-within year_.

|  | *Minimal* |  | *Nestbox* |  | *Spatial proximity* |  | *Breeding enviro. sim.* |  |
| --- | --- | --- | --- | --- | --- | --- | --- | --- |
| | Est(SE) | Rel to $V_P$ | Est(SE) | Rel to $V_P$ | Est(SE) | Rel to $V_P$ | Est(SE) | Rel to $V_P$ |
| **Laying date** |  |  |  |  |  |  |  |  |
| $V_{By}$ | 49.082 (8.855) | 0.600 (0.108) | 49.018 (8.842) | 0.602 (0.109) | 49.115 (8.907) | 0.490 (0.088) | 49.045 (0.848) | 0.046 (0.008) |
| $V_A$ | 6.566 (0.849) | 0.080 (0.010) | 6.271 (0.794) | 0.077 (0.010) | 6.100 (0.753) | 0.061 (0.008) | 5.448 (0.758) | 0.054 (0.008) |
| $V_{Pe}$ | 6.560 (0.864) | 0.080 (0.011) | 4.866 (0.808) | 0.060 (0.010) | 3.797 (0.764) | 0.038 (0.008) | 4.618 (0.799) | 0.046 (0.008) |
| $V_{Nb}$ | - | - | 2.426 (0.233) | 0.030 (0.003) | - | - | - | - |
| $V_{SPATIAL}$ | - | - | - | - | 21.663 (4.022) | 0.216 (0.040) | - | - |
| $V_{BREED\_ENV}$ | - | - | - | - | - | - | 21.960 (3.736) | 0.218 (0.037) |
| $V_R$ | 19.570 (0.422) | 0.239 (0.005) | 18.829 (0.420) | 0.231 (0.005) | 19.505 (0.414) | 0.195 (0.004) | 19.487 (0.416) | 0.194 (0.004) |
| $V_P$ | 81.778 | - | 81.409 | - | 100.180 | - | 100.559 | - |
| $V_{P-within\ year}$ | 32.696 | - | 32.391 | - | 51.065 | - | 51.514 | - |
| $h^2$ | **0.201 (0.058)** | - | **0.194 (0.054)** | - | **0.119 (0.026)** | - | **0.106 (0.010)** | - |
| **Hatching date** |  |  |  |  |  |  |  |  |
| $V_{By}$ | 46.095 (8.306) | 0.648 (0.117) | 45.903 (8.278) | 0.650 (0.117) | 45.924 (8.280) | 0.544 (0.098) | 45.773 (8.255) | 0.520 (0.094) |
| $V_A$ | 5.163 (0.674) | 0.073 (0.009) | 4.876 (0.621) | 0.069 (0.009) | 4.641 (0.580) | 0.055 (0.007) | 4.184 (0.585) | 0.048 (0.007) |
| $V_{Pe}$ | 4.840 (0.682) | 0.068 (0.010) | 3.188 (0.627) | 0.045 (0.009) | 2.418 (0.585) | 0.029 (0.007) | 2.907 (0.614) | 0.033 (0.007) |
| $V_{Nb}$ | - | - | 2.247 (0.202) | 0.032 (0.003) | - | - | - | - |
| $V_{SPATIAL}$ | - | - | - | - | 16.481 (3.013) | 0.195 (0.036) | - | - |
| $V_{BREED\_ENV}$ | - | - | - | - | - | - | 20.231 (3.241) | 0.230 (0.037) |
| $V_R$ | 15.014 (0.338) | 0.211 (0.005) | 14.430 (0.336) | 0.204 (0.005) | 15.007 (0.332) | 0.178 (0.004) | 14.979 (0.333) | 0.170 (0.004) |
| $V_P$ | 71.113 | - | 70.082 | - | 84.471 | - | 88.072 | - |
| $V_{P-within\ year}$ | 25.018 | - | 24.179 | - | 38.547 | - | 42.299 | - |
| $h^2$ | **0.206 (0.051)** | - | **0.197 (0.047)** | - | **0.120 (0.024)** | - | **0.099 (0.020)** | - |

**Fig. 1.**
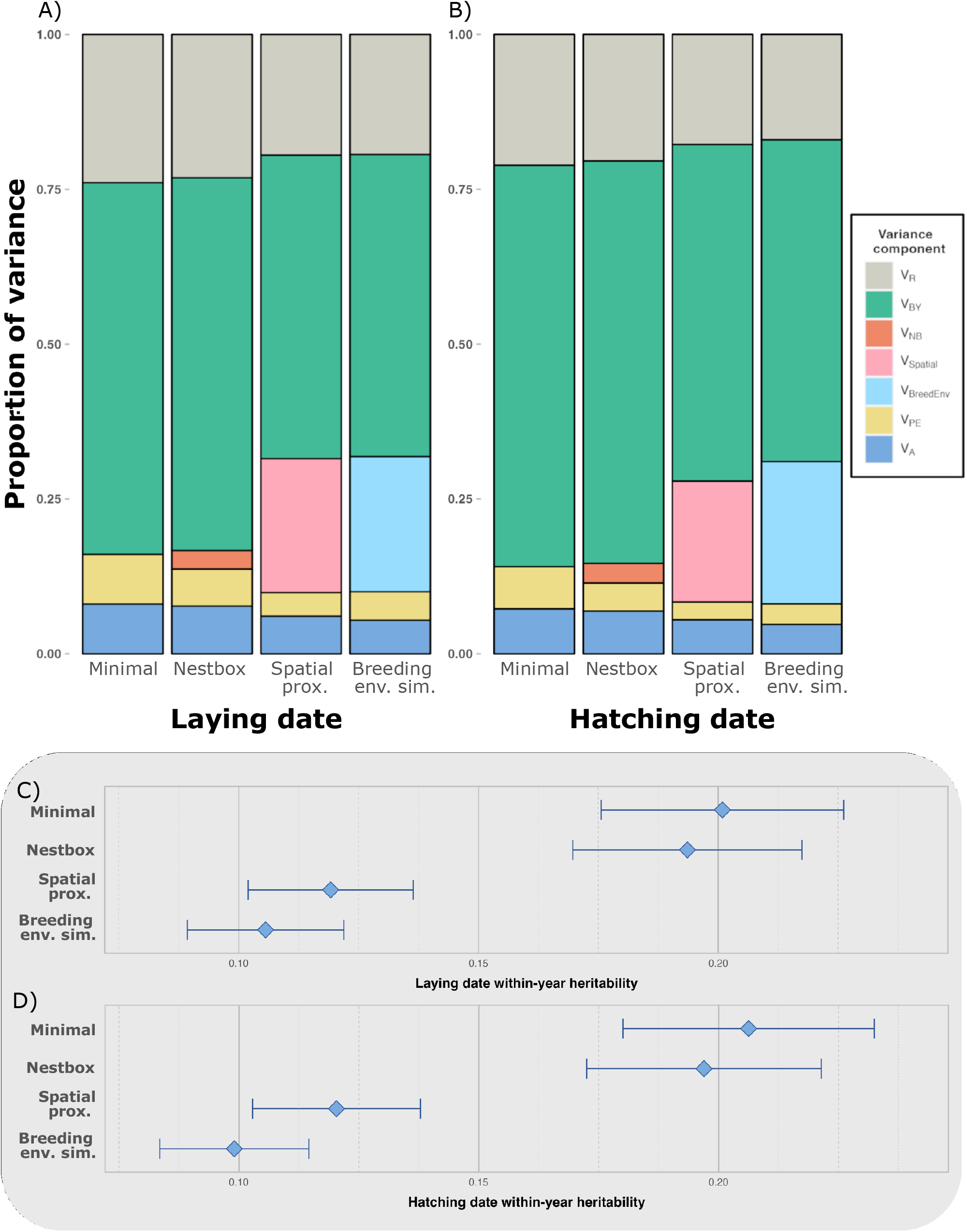
Proportion of variance assigned to each component for the 4 different models (specific information on all models in the methods), for (A) laying date and (B) hatching date. Colours correspond to each component. *V*_BY_ = breeding year, *V*_SPATIAL_ = spatial proximity matrix, *V*ENVIRONMENT = environmental similarity matrix, *V*_PE_ = focal individual permanent environment effect, *V*_NB_ = nest box random effect, *V*_A_ = focal individual additive genetic effect, *V*_R_ = residual variance C) and D) show within-year heritability estimates (estimated as the proportion of variance explained by *V*_A_ out of the within year *V*_P_, for all 4 models for laying date (C) and hatching date (D). Error bars show standard error.

Both the spatial proximity and breeding environment similarity terms significantly improved the model fit compared to the nestbox model (nestbox model vs spatial model: χ2 = 523, p < 0.001, nestbox model vs breeding environment model: χ2 = 259, p < 0.001)). The spatial model was a significantly better fit to the data than the breeding environment model (χ2 = 264, p < 0.001).

The spatial proximity matrix and the breeding environment similarity matrix explained 21.7 *±* 4.0% and 21.9 *±* 3.7% of the variation seen in laying date, respectively. The incorporation of these S-matrices reduced the proportion of variation explained by breeding year by 19% and 16% compared to the nestbox model (from 60.2% in the nestbox model to 49.0% in the spatial model and 48.8% in the breeding environment model). Both models produced similar results for the proportion of variation explained by additive genetic effect (i.e. in relation to narrow-sense heritability (spatial 6.1% and breeding environment, 5.4%). The individual permanent environment effect was also similar, breeding environment model compared to the spatial model (3.8% vs 4.6%). The amount of residual variation was reduced when including either matrix, by around 16% (from 23.1% to 19.4% and 19.3%).

Including either S-matrix led to within-year heritability estimates of approximately two thirds the estimate from the nest-box model. The spatial model estimated a narrow-sense heritability of 11.9 *±* 1.7% and the breeding environment model was slightly lower at 10.6 *±* 1.6%, reducing the heritability estimate by over 40% (Table 1 and Figure 1C).

### Hatching date

Laying date and hatching date are very closely correlated (Pearsons *r*(10893) = 0.952, *r*^2^ = 0.905, interval between laying and hatching has a median of 21 days with a SD = 2.735), so we therefore expected very similar results. As for laying date, breeding year explained the greatest proportion of variation in hatching date in all four models (ranging from 52.0 *±* 9.4% to 65.0 *±* 11.7%: Table 1 and Figure 1B). Compared to each equivalent laying date model, breeding year contributed just under 5 percentage points more to variation in hatching date. Including nest box as a random effect significantly improved the model fit compared to the minimal model (χ2 = 288, p < 0.001), with the nest box random effect explaining 3.2 *±* 0.3% of the variation. Within-year heritability only decreased slightly, from 20.6 *±* 2.6% in the minimal model, to 19.7 *±* 2.4% in the nestbox model (Table 1 and Figure 1D). The effect of mothers age at breeding does not change much across models, but is slightly lower for hatching date than for laying date (Supplementary Table 5)

Both S-matrices significantly improved the model fit compared to the nestbox model (nestbox model vs spatial model: χ2 = 583, p < 0.001, nestbox model vs breeding environment model: χ2 = 313, p < 0.001). As with laying date, the spatial model was a significantly better fit than the breeding environment model (χ2 = 270, p < 0.001).

The spatial proximity matrix explained 19.5 *±* 3.6% of variation seen in hatching date, and the breeding environment similarity matrix explained 23.0 *±* 3.7%. Incorporation of either S-matrix reduced the proportion of variation explained by breeding year compared to the nestbox model - from 65.0% to 54.4% (spatial) and 52.0% (breeding environment).

There was little difference between the proportion of variance explained by additive genetic effects between the spatial and breeding environment model (spatial model, 5.5% and breeding model, 4.8%). The individual permanent environment effect explained was also very similar between the spatial model and the breeding environment model (2.9% vs 3.3% respectively).

The proportion of residual variance very similar same between models (17.8% *±* 0.4% and 17.0% *±* 0.4%). Including either matrix reduced the within-year heritability estimates by 39% and 50% compared to the nestbox model (19.7% to 12.0% and 9.9%: Table 1 and Figure 1D).

## Discussion

We demonstrate the importance of accounting for small-scale environmental variation when estimating heritability of a phenological trait in a wild population, showing that neglecting this variation leads to overestimation of heritability. The heritability of breeding timing in this population of great tits was almost halved when the similarities in the breeding environments experienced by individuals were taken into consideration. Previous assessment of the effects of spatial and environmental similarity on phenological timing and heritability have been limited: a single previous study used mother-daughter regressions across three distance classes of female natal dispersal, and observed decreasing heritability over greater dispersal distances (H. P. Van Der Jeugd and McCleery, 2002). Our study advances understanding in this area by using the full pedigree within multi-matrix animal models, and accounting for smaller-scale environmental similarity in a comprehensive way.

Similar results were found for both laying date and hatching date, which is expected as they are closely related. We found larger effects of year, and slightly reduced additive and permanent environment effects when considering hatching date compared to laying date. This may be due to the fact that individuals are able to adjust their hatching date after laying, by controlling the number of eggs laid, and the onset of incubation (Cresswell and McCleery, 2003; Simmonds et al., 2017). This could allow hatching to vary more between years in response to the environment, however the difference is small, and does not result in significant differences in heritability estimates between the two traits.

We also found that the relative size of the permanent environment effect was reduced for these phenological traits when including either matrix compared to a random nestbox effect, suggesting that some variation previously assigned to within-individual variation was better explained by the breeding environments those individuals experience across their lifetime. Hence, if breeding location was not accounted for, analyses might incorrectly assign variance to the female rather than to the environment (i.e. a bird laying earlier in a “good” location may be assumed to be early due to an inherent property of the individual rather than the site per se).

In the context of considering spatial proximity and also environmental similarity, we found that there was relatively little correlation between these two factors in our population (mantel correlation of 0.192). Further, both matrices separately accounted for approximately 20% of the variation observed in laying date and in hatching date, suggesting that they capture separate but similarly-sized effects. Further work involving direct comparison of both spatial and environmental similarity in this population, and other populations, would be very valuable, but currently challenging given the methodological difficulties in estimating variance components simultaneously. However, simulation studies, where the precise environmental structure and data structure can be controlled may be particularly fruitful in gaining further insights on where spatial or environmental sources of similarity will be particularly important sources of individual phenotypic variation.

One of the reasons for the previous lack of consideration in environmental similarity in shaping heritability estimates may be the challenges associated with measuring this. Indeed, considering environmental similarity is limited by the choice of environmental factors used to create a measure of environmental similarity. All such studies have to make assumptions about how to choose factors which may represent the relevant environment. In doing so, we may not capture the full importance of the environment, and methods which combine various factors of the environment (such as ours here) may be masking some of their effect. While it is difficult to fully mitigate this issue (due to physical limitations on data collection, and complexities with overloading models given how multi-dimensional the environment is), long term study systems with detailed information (such as ours) provide a first step to integrating this consideration properly when considering heritability in the wild.

The causes of any genotype-environment covariance are also important. If it results from a genetically mediated breeding environment choice then this could actually be considered part of the ‘genetic’ heritability of the trait. Although consistent differences between wild individuals in their habitat choice are commonplace (Bell et al., 2009; Leclerc et al., 2016) few studies have been able to quantify the degree to which variation in habitat choice is driven by genetics (but see Gaither et al., 2018; Jaenike and Holt, 1991). In cases where breeding habitat choice is, at least partly, genetically determined, and thus breeding environments are, to some degree, heritable, the overall heritability or effective heritability, would be underestimated by assuming similarity in the environments used by individuals is driven purely by non-genetic processes (e.g., limited dispersal). For example, genetic heritability of habitat selection has recently been found in homerange habitat composition in a roe deer population, incorrectly downward biasing heritability estimates of behavioural and morphological traits (Gervais et al., 2022) It is currently unclear how heritable environmental choice is in birds, and whether it is a mechanism through which genetics influence lay date. Thus, our findings may represent only a part of the underlying mechanism.

Our primary results align well with current general investigation into this area, with other studies showing the importance of accounting for spatial variation when estimating heritability with quantitative genetic models (Regan et al., 2017; Rutschmann et al., 2020; Stopher et al., 2012; Thomson et al., 2018; Henk P. Van Der Jeugd and McCleery, 2002). Birds that breed seasonally can be influenced by the environment at small local scales, and the timing of laying of great tits has been shown to vary with food availability and quality (Cole et al., 2021; Hinks et al., 2015), habitat composition (Matthysen et al., 2021) and territory size (Wilkin et al., 2006). Indeed, phenotypic plasticity is important for short term adjustments, and in the Wytham population individual plastic adjustments in timing of breeding have been key in tracking the rapidly changing environment (Charmantier et al., 2008a). The impact of these small-scale environmental effects on estimates of the heritability of timing of breeding in wild bird populations has not previously been quantified. Furthermore, the reduction in permanent environment effects when considering environmental similarity that we report here has also been found in other systems, such as for behavioural traits in roe and red deer (Gervais et al., 2022; Stopher et al., 2012).

Our analyses clearly highlight the need for more careful consideration of breeding environment similarity across space, especially in wild populations where the environment can vary greatly. Specifically, as we found that breeding environment similarity between individuals explains similar amounts of variation as spatial proximity, it may generally be that in some cases it is perhaps desirable to use this measure of breeding environment similarity instead of spatial proximity to capture variation in phenotypes due to both space and environment. Yet, how the relative proportion of variation explained by spatial compared with breeding environment similarity matrices varies depending on context remains unknown, as well as how life history of the species may shape this. For example, generation time and natal dispersal distance will impact how likely related individuals are to experience similar environments. In species with long dispersal distances we may expect environmental similarity to play a more important role in contribution to variation in a trait; in contrast, for a species with limited dispersal, or even inherited territories, it would be less important to consider this. As such, investigations around when it is most appropriate to consider environmental similarity over spatial similarity would now be useful. Such analyses would likely focus on considering factors such as the dispersal distances of a species and the grain of habitat variability experienced by the population. For example, in species with shorter dispersal distances and a larger grain of habitat variability (meaning environments close by are more similar), offspring are more likely to experience similar breeding environments to their parents. However, if the grain of habitat variability is very small (meaning nearby environments could be quite different), offspring that disperse short distances may end up actually experiencing different environments to their parents. In such cases, it becomes more important to consider not just spatial proximity but also breeding environment similarity. This could potentially be explored in future research, to gain a better understanding of what measurements of common environment effects are most appropriate for a given population or species.

In a broader sense still, the possibility of inherited environments raises questions about how selection of environments could be transmitted from parents to offspring. It is important to consider the heritability of environment choice so that heritability estimates are not biased downwards by removing genetic variation that underpins similarity between parents and offspring. In this context, our study population could be used to investigate whether individuals actively choose to nest in environments more similar to where they were born, which could be contributing to the genetic heritability of the trait.

### Summary

Understanding the additive genetic and environmental contributions to phenotypic variation in phenological traits is important for a range of questions about their evolution, and for understanding their potential to respond to changing environments. It is clear that if common environment effects are not considered, estimates of heritability and trait evolvability will be biased. Our study shows that accounting for the shared environment is important for understanding the genetic basis of reproductive timing variation in wild individuals, and is useful for enabling understanding of the causes and consequences of different components of phenotypic variation. As global change continues to impact phenology, it is crucial to continue to develop methods that account for small-scale environmental variation. Our approach, which includes a measure of breeding environment similarity, aims to capture both spatial autocorrelation and environmental similarity of individuals not necessarily close in space. Long-term study systems which detail fine-scale individual level information across generations (such as ours) provide great opportunity for considering space and environment types, and would therefore will be useful in examining the effects of global change within a natural population in this context.

## Methods

### Study System

This study used data from the long-term study of great tits (*Parus major*) in Wytham Woods, Oxfordshire over the years 1960 to 2022. Great tits rely on timing their breeding with the local temporally variable food supply, timing peak resource requirements of nestlings with a peak in their primary consumer prey, which in turn depends on the leaf development of deciduous trees (Hinks et al., 2015). Great tits are cavity nesters and in this population the majority of the breeding population nests in the 1019 nest boxes placed throughout the woods (Harvey et al., 1979), which are monitored over the breeding season (from March – May) each year following a standardised protocol (Perrins et al., 1965).

Birds are uniquely identified with metal BTO (British Trust for Ornithology) rings; since 2007 all birds have also been fitted with plastic rings which contain a passive integrated transponder (PIT tag). Parent birds were identified at the nest when provisioning young. Radio Frequency Identification (RFID) antenna can read the PIT tags, so are placed around nest box entrance holes, allowing identification of individuals whilst feeding nestlings, without the need to catch them. Birds that cannot be identified using this method (likely unringed individuals) are trapped at the nest box when nestlings are at least 10 days old. They are then fitted with metal BTO leg rings and PIT tags. All individual identification was done by catching before the use of PIT tags. All nestlings are ringed and PIT tagged on day 15 before they leave the nest. For nests that fail or are abandoned before fledging often only mother ID is recorded. Mist netting is carried out over the autumn and winter to catch and ring as many immigrant birds as possible.

### Breeding data

Two traits were used to represent breeding timing: laying date, defined as the day the first egg of a clutch was laid, assuming that females lay one egg a day early in the morning, and hatching date, defined as the day the first egg was hatched. Nest boxes are visited at least once a week from late March, until eggs are found; if there is more than one egg on first observation of eggs in the nest, the date of first egg is inferred by counting back assuming one egg is laid per day. Once eggs are observed to be warm, indicating incubation has begun, nests are not be visited again until the expected hatch date (12 days after clutch completion). Onset of incubation and incubation duration can vary between individuals; if nestlings are not observed on the predicted hatch date, the nests are visited every other day until hatching or until the nest is declared abandoned. Newly hatched nestlings have a distinctive appearance, allowing fieldworkers to establish whether the largest young in a nest are more than a day old. If there any ambiguity, 3-4 of the largest nestlings are weighed to determine age (Supplementary Table 3). This protocol ensures that all hatch dates should be accurate to *±*1*day*.*DatesareexpressedhereinAprilDays*(1*stApril* = 1).

Overall there were 17,996 recorded breeding attempts over 63 years, from 1960 to 2022. From this sample only breeding attempts where the mother was identified and had a recorded laying date were kept (removing 5,848 attempts); further, any breeding attempts from known second broods were removed (260 and 19 attempts for mothers and known fathers respectively), as well as removing any laying dates that were 30 days after the first 5% of laying dates that year within subsets of the woods (the study site is split into 9 sections, based loosely on habitat types, for logistic purposes; 126 attempts), to account for unknown second broods (e.g. Van Der Jeugd and McCleery, 2002), and any broods that were experimental manipulated or do not have complete habitat data (1,673 attempts); overall this left a sample of 11,658 breeding attempts for analysis. We kept breeding attempts where mothers were known, in preference to fathers, as here we opt to treat laying date as a maternal trait both given that the heritability of laying date in males is less than a fifth that of females (Evans et al., 2020) and because this makes the quantification of shared breeding environments tractable.

### Pedigree construction

Identification of individuals at the nest box enabled the creation of a social pedigree across 62 years. The social pedigree assumes that the adult birds identified incubating or feeding nestlings at a nest are the biological parents. Other than clerical error, the maternal pedigree should be accurate as there are no known cases of maternal identity mismatching social parent identity (Patrick et al., 2012). There are relatively low levels of extra-pair paternity recorded in this population (of the order of 12%: Patrick et al. 2012). Extra-pair paternity at this level is not thought significantly influence quantitative genetic estimates assuming it is not strongly biased with respect to traits of interest (Charmantier and Réale, 2005; Firth et al., 2015), and given our focus on timing as a maternal trait we consider this a reasonable assumption. The pedigree analysed here includes 14,506 individuals (including only individuals with recorded breeding attempts who contribute information to this analysis) and extends for up to 35 generations, with 7,431 maternities and 6,761 paternities, 4,260 full siblings, 3,033 maternal half siblings and 2,114 paternal half siblings.

### Analysis

We constructed animal models in ASReml-R (Butler et al., 2007) to partition the phenotypic variance in each of the traits into genetic and environmental variance components and reassess the heritability (Kruuk, 2004; Wilson et al., 2010). The pedigree was used to create a matrix of expected relatedness between all individuals, allowing the consideration of many different relationships instead of just parents and offspring. Raw laying/hatching date data was used, given in April Days (April 1st = 1). The age of the female at breeding was included as a fixed effect as a 2-level factor, first-year breeders (1-year-old) or older adult (older than 1 year) (Evans et al., 2020).

In all models we included year of breeding as a random effect (*V*_BY_) to partition the variance attributable to variation in the environment during the year of breeding, in this population previous studies have found major phenotypic plasticity across years (Charmantier et al., 2008). We also included individual identity of the breeding female, linked to the pedigree, as a random effect (*V*_A_) to estimate the additive genetic effect, which is the influence of the genes that belong to the individual in which the trait was measured. The individual identity was also included as a permanent environment effect (*V*_PE_), to adjust for multiple records of individuals over years, accounting for non-heritable effects that will cause variation that is conserved across the repeated records of individuals (e.g. natal effects) (Kruuk and Hadfield, 2007; Lynch and Walsh, 1998).

### Accounting for environmental similarity

First, a model was run with just the factors outlined above (the minimal model; model 1). We then ran additional models, extending the minimal model by adding: an individual nest box random effect (model 2), a matrix of spatial proximity (model 3), a matrix of environmental similarity (model 4), and a model including all three simultaneously.

Model 1 (minimal model) simply decomposes the phenotypic variance (*V*_P_) into breeding year effect (*V*_BY_)), female permanent environment effect (*V*_PE_), genetic effects (additive genetic effect of female (*V*_A_), and the residual variance (*V*_R_) which accounts for variation arising from environmental effects that have not been explicitly included in the model.

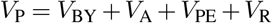

Model 2 (nestbox model) included “nest box” as a random effect (*V*_NB_), which accounted for similarities in breeding timing of different females breeding in the same boxes over time that was due to similar breeding environments. Individual nest boxes were used between 1 and 35 times each.

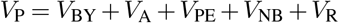

Models 3 and 4 include an ‘S-matrix’ that describes the similarity of a non-genetic effect between individuals, and works in the same way the genetic relatedness matrix does, to estimate the contribution of variance associated with environmental effect. This was first applied by Stopher et al., (2012). Model 3 (spatial proximity model) contained a spatial proximity matrix. This was constructed by taking the breeding location of each individual bird and calculating the distance between all possible combinations of birds across all years. If an individual was recorded breeding more than once in different nest boxes (28% of individuals), the mean location point was taken. We expect this will not affect the results significantly as breeding dispersal in this population is limited to short distances: median of 60.75 metres (Supplementary Figure 3).

The distance values were scaled from 0 to 1, with 1 along the diagonal such that individuals have a similarity of 1 with themselves, and 0 was the maximum distance between a pair of individuals (3971.92m). The matrix was linked to the animal model to estimate the proportion of variance explained by the distance between individuals (*V*_SPATIAL_):

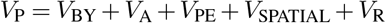

Model 4 (breeding environment model) was constructed to account for effects of small-scale environmental variation, independently of the effect of distance. This was done by including a matrix of breeding environment similarity between individuals. We included the following factors: altitude, edge distance index, northness, oak-richness within 75m, and population density (expressed as the square root of territory size).

In this population of great tits, females lay earlier at lower altitudes, on north facing slopes, at more interior sites, when oak tree density within 75m of the box is higher, and at lower population densities (when they have larger territories) (Wilkin et al., 2007b, 2007a, 2006).

The environmental factors were chosen as they are factors that vary over small spatial scales in Wytham, and have been previously shown to be related to variation in laying date, are likely to have remained the same over the years of the study (i.e. physical features of the environment and not climatic factors; boxes have a fixed location).

As with distance, for birds that were recorded breeding more than once, a mean for all environmental values over years was taken. Breeding dispersal is minimal (median = 60.75 metres: Supplementary Figure 3) and therefore measures of breeding environment between boxes used for consecutive breeding attempts are highly repeatable. We then used methods suggested by Thomson et al. (2018) to combine the environmental measures to values of breeding environment similarity between all individuals. Each variable was centred and scaled, and then combined using Euclidean distance measure in multivariate space between all individuals with every other individual, to obtain the straight-line distance between 2 vectors of environmental measures in multivariate space. This creates a similarity matrix which aims to capture a substantial amount of the similarity in the environment experienced by individuals as a single value. This similarity value was again scaled to give a value of 1 along the diagonal, with 0 as the distance between birds in the most dissimilar environments (histogram of the distribution of breeding environment similarity values in Supplementary Figure 4).

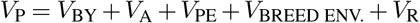

Finally, we attempted to run a model including both the spatial proximity and breeding environment similarity matrices in one model, however we encountered convergence problems with the model, likely due to a lack of power in the data to decompose both environmental matrix effects.

In order to understand the relationships between genetic relatedness, spatial proximity, and environmental similarity, we carried out two sets of examinations. First, we visualised the relationship between the two matrices: spatial proximity, and environmental similarity, to ensure there was representation of genetically related individuals experiencing more and less close/similar environments. Second, we also ran mantel correlation tests to broadly quantify the correlations between the different matrices (Mantel, 1967).

### Assessing heritability

We used the within-year phenotypic variance, which is the sum of all variance effects except breeding year, conditioned upon the fixed effect of female age at breeding, to estimate the heritability (Evans, Postma and Sheldon, 2020). This is for two reasons: firstly, when selection is estimated for these traits it is typically done on a year-specific basis (Noordwijk, McCleery and Perrins, 1995; Charmantier et al., 2008). Secondly, there has been a long-term advancement in breeding timing in this population; as most individuals in the population only live to breed for 1 or 2 years, this will likely lead to overestimation of annual variance above what will be actually experienced by individuals.

Heritability is therefore given as the proportion of within-year phenotypic variation (*V*_Pwithin-year_) assigned to additive genetic variance (*V*_A_).

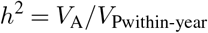

The proportion of variance explained by each of the variance components was calculated as the ratio of the relevant component to the within-year phenotypic variance *V*_Pwithin-year_. To assess the significance of the random effects, we used likelihood ratio tests, assuming a χ2 distribution with one degree of freedom (Wilson et al., 2010).

## Acknowledgements

We thank all Wytham fieldworkers who have worked on the tit study over many years. Work funded by the Edward Grey Institute for Field Ornithology, and the long-term population study has been supported by numerous funding sources, including recently by grants from BB-SRC (BB/L006081/1), ERC (AdG250164), and NERC (NE/K006274/1, NE/S010335/1). Authors have no conflict of interest to declare.

## Author contributions

All authors conceptualised the idea. All authors have participated in data collection. Carys V. Jones conducted data analysis with input from Charlotte E. Regan. Carys V. Jones produced first draft of the manuscript. Charlotte E. Regan, Ella F. Cole, Josh A. Firth, and Ben C. Sheldon provided detailed feedback on methodology, and contributed critically to drafts.

## Data and code availability

Code and data to reproduce all analyses is available at https://github.com/carysvjones/AnimalMod_Envir.git

