## Supplementary Info for "Environmental similarity between relatives reduces heritability of reproductive timing in wild great tits"

### **Section 1 - Regression analysis**

Repeating analysis from Van der Jeugd & McCleery 2002 with additional data.

Parent-offspring regressions use phenotypic similarity between related individuals to estimate heritability (Lynch and Walsh, 1998). For single parent-offspring regression, the coefficient estimate from a linear regression of mother’s mean phenotype against daughter’s mean phenotype is multiplied by the inverse proportion of expected genes shared to estimate the heritability; hence in this case it is multiplied by two as the offspring share, on average, half their genes with the parent (Boag and van Noordwijk, 1987). The parent-offspring regressions were calculated for laying and hatching date, and then based on previous analysis of this population by (Van Der Jeugd and McCleery, 2002) the analysis was repeated by forming groups of relatives with different dispersal distances, to emphasis the effect of shared environments. We present this updating of the initial analysis of van der Jeugd and McCleery (2002) for two reasons: (1) It was an early attempt to estimate formally the effect of shared environments on heritability of timing and we consider it interesting to repeat with a substantially larger data set; (2) it provides a simple visual interpretation of the effect of increased spatial distance on heritability.

#### Parent-offspring regressions

Parent-offspring regressions use phenotypic similarity between related individuals to estimate heritability (Lynch and Walsh, 1998). Using just female birds, from the pedigree there were 3,371 relationships where both a daughter and mother had recorded first egg lay dates; 95% (N = 3,187) of those pairings also had a recorded hatch date (the decrease in number of hatch dates compared to laying dates is due to abandoned nests before or during incubation). Overall, this includes 4,604 unique birds (some daughters will also be included as mothers, and some mothers will have multiple daughters). Due to significant differences in mean laying and hatching date between years and age classes all dates were standardised by year and age. The age of all birds was known to good degree of certainty, either from records of their birth, or from aging from plumage when caught. If an individual had multiple recorded laying or hatching dates (46% of individuals included in this study) their lifetime mean was taken. If a mother had multiple recorded daughters who bred, the mother was included once for each mother-daughter pairing.

##### *Accounting for dispersal distance*

Natal dispersal distance was defined as the distance moved by an individual between their natal box (the nest box they were born in), to the box where they first breed (most commonly at age 1). The median natal dispersal for females was 786m, with 62% breeding within 1km of where they were born (Supplementary Figure 1). Natal dispersal distance is a straightforward way to begin to account for the shared environment between mothers and daughters. The environment in Wytham is heterogenous over small scales; for example across the 385ha woodland altitude varies by 100m (Wilkin, Perrins and Sheldon, 2007). So, we may expect offspring that disperse shorter distances to end up in a more similar environment to their parents.

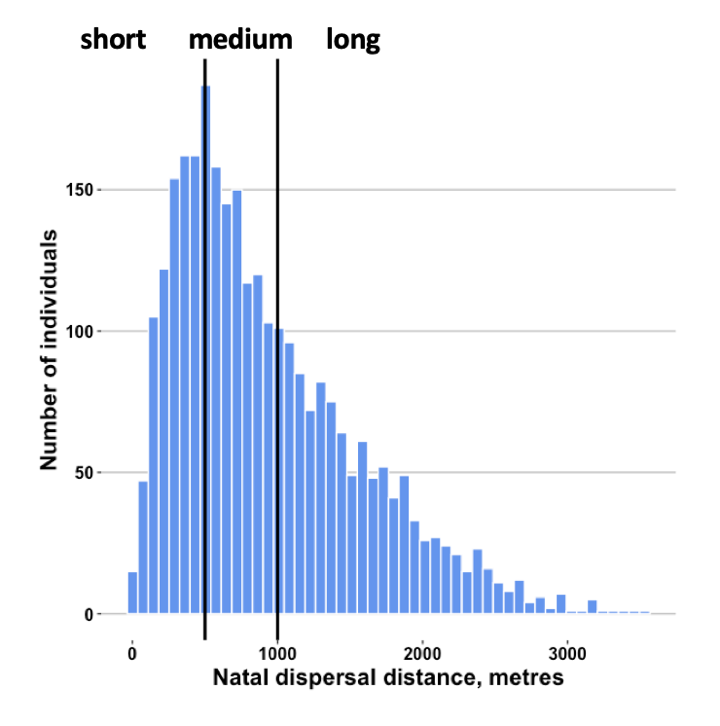

**Supplementary Figure 1 – Natal dispersal distance** – Natal dispersal distance of females in population. Vertical black lines show where the data was split to create the short, medium and long dispersal groups, labelled above the plot. Further details about dispersal groupings in Table 1, n = 3371.

Natal dispersal distance was calculated for all daughters. To replicate the analysis from Van Der Jeugd and McCleery (2002) we then split the mother-daughter pairings into three distance groups: ‘short’ (0-500m), ‘medium’ (500-1000m) and ‘long’ (>1000m) dispersers. The number of pairs and mean dispersal distance within each group are shown in Supplementary Table 1, and a histogram showing the range in dispersal distances overall, with lines indicating where the groups are split, is shown in Supplementary Figure 1 – note that the groupings are artificial, but that these distance classes do split the data into three approximately equal sized groups. The parent-offspring regressions were recalculated for each of these dispersal groupings, and heritability estimated for each.

**Supplementary Table 1 – Dispersal distance groupings *–*** The data were split into 3 groups, short, medium and long dispersers, depending on the distance daughters moved from their natal to first breeding box. The table shows the number of mother-daughter pairings in each group (N), and the distance range and mean distance dispersed within each group.

| ***Trait & Group*** | ***N*** | ***Distance range*** | ***Median distance*** |
| --- | --- | --- | --- |
| **Laying Date** |  |  |  |
| short | 986 | 0-500m | 315m |
| medium | 1095 | 500-1000m | 718m |
| long | 1290 | >1000m | 1480m |
| **Hatching Date** |  |  |  |
| short | 946 | 0-500m | 318m |
| medium | 1032 | 500-1000m | 720m |
| long | 1209 | >1000m | 1482m |

#### Regressions results

Results are shown in Supplementary Table 2, alongside the estimates of heritability for laying date from Van Der Jeugd and McCleery (2002).

**Supplementary Table 2 – Regression model output** *–* _(1)_ is results from (Van Der Jeugd and McCleery, 2002) and _(2)_ are from this study. Hatching date vs laying date. h^2^ is the within-year heritability, as the ratio of V_A_ to V_P-within years_, SE is the standard error of heritability. N is the number of pairings of mothers and daughters used in each analysis.

|  | ***short*** | | | ***medium*** | | | ***long*** | | | ***overall*** | | |
| --- | --- | --- | --- | --- | --- | --- | --- | --- | --- | --- | --- | --- |
|  | *h^2^* | *SE* | *N* | *h^2^* | *SE* | *N* | *h^2^* | *SE* | *N* | *h^2^* | *SE* | *N* |
| Laying date_(1)_ | 0.40 | 0.10 | 399 | 0.25 | 0.09 | 458 | 0.07 | 0.10 | 476 | 0.24 | 0.06 | 1332 |
| Laying date_(2)_ | 0.37 | 0.07 | 986 | 0.20 | 0.07 | 1095 | 0.15 | 0.06 | 1290 | 0.23 | 0.04 | 3371 |
| Hatching date_(2)_ | 0.35 | 0.07 | 946 | 0.18 | 0.07 | 1032 | 0.16 | 0.04 | 1209 | 0.24 | 0.04 | 3187 |

#### Laying date

The regressions using all data gave a heritability estimate of 23 ± 4% (Supplementary Figure 2a). When data was split by dispersal distance groups, there was a 2.5-fold difference in heritability estimates between short and long dispersers (Supplementary Figure 2b). The analysis of short dispersers gives a heritability estimate of 37 ± 7%, whilst the analysis of long dispersers yielded an estimate of 15 ± 6%.

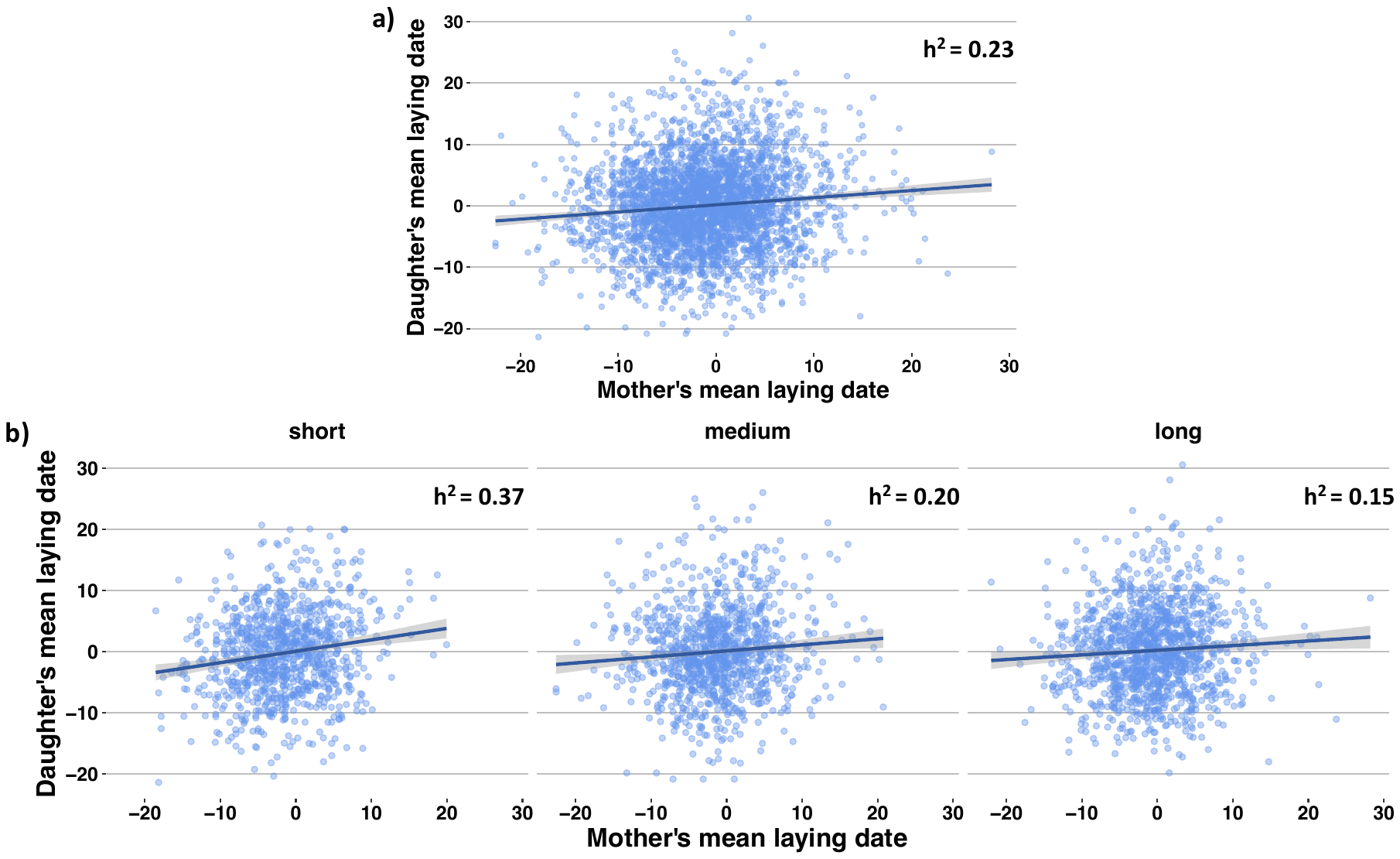

**Supplementary Figure 2 – Heritability split by dispersal**. Simple linear parent-offspring regressions for laying date. a) regression with all individuals, b) shows regressions of 3 dispersal distance groups, short, medium and long. h^2^ is heritability calculated as twice the estimates coefficient, in the case of single-parent offspring regressions.

#### Hatching date

Heritability estimates for hatching date were strikingly similar to those for laying date. The overall parent-offspring regression analysis yield an estimated heritability of hatching date as 24 ± 4%. When the data was split up the short disperser’s heritability was estimated to be 2.2 times that of the long dispersers’ heritability (35 ± 7% compared to 16 ± 4%).

#### Summary

It is striking that the heritability estimates calculated using parent-offspring regressions with the updated data are very similar results to those found in earlier estimates by Van der Jeugd and McCleery (2002), despite being estimated with more than twice the quantity of data. Heritability estimates declined with increasing dispersal distance of daughters, clearly demonstrating the effect of a common environment on similarity in timing of breeding, with offspring that nest closer to where their parents did also breeding at a more similar time. However, we know parent-offspring regressions do not utilise the full extent of data we have available and often overestimate heritability. In our analysis, heritability estimated using the animal model gave lower estimates in all models compared to the overall parent-offspring regressions, agreeing with previous work (van Noordwijk, 1984; Merilä and Sheldon, 2000; Sheldon, Kruuk and Merilä, 2003; Kruuk and Hadfield, 2007; Postma and Charmantier, 2007; Evans, Postma and Sheldon, 2020).

### **Section 2 - Methods and Results**

**Supplementary Table 3 – Chick hatch weights** - When unsure of hatch day, chicks can be weighed to estimate day of hatching. Newly hatched chicks are clearly identifiable by eye, but chicks slightly older are more difficult to judge. Weighing 2 or 3 of the largest chicks and taking an average weight can help estimate the day of hatching using the table below.

| **Day** | **Great tit** |
| --- | --- |
| 1 (hatch day) | 0-2g |
| 2 | 2-3 |
| 3 | 3-4.3 |
| 4 | 4.3-6.0 |
| 5 | 6.0-7.7 |
| 6 | 7.7-9.4 |
| 7 | 9.4-11.2 |
| 8 | 11.2-13.0 |
| 9 | 13.0-14.6 |
| 10 | 14.6+ |

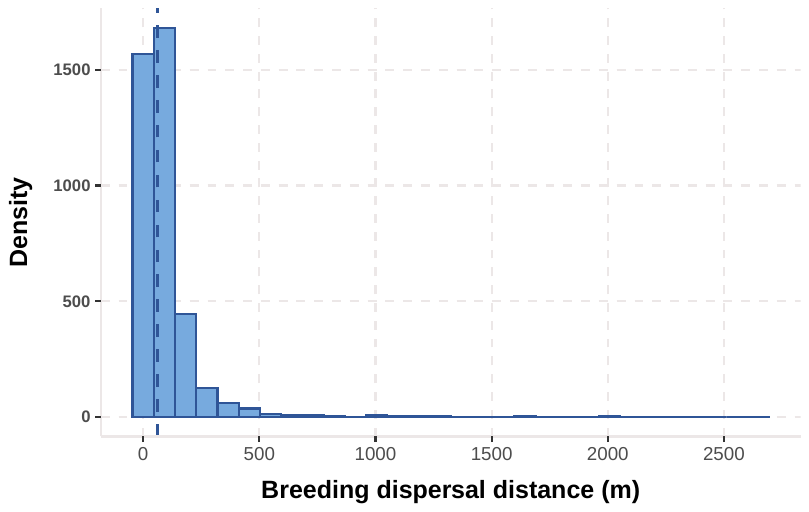

**Supplementary Figure 3 – Breeding dispersal.** Histogram of the distances moved in metres between nest boxes by females who were recorded breeding in multiple years (n = 2589 females, 954 females were recorded breeding in more than 2 years (max 8 years), and 958 (14.6%) breeding dispersal movements were 0m i.e. a female re-nesting in the same box in consecutive years). Breeding dispersal has a median of 60.75m, considerably less than natal dispersal which has a median of 786m.

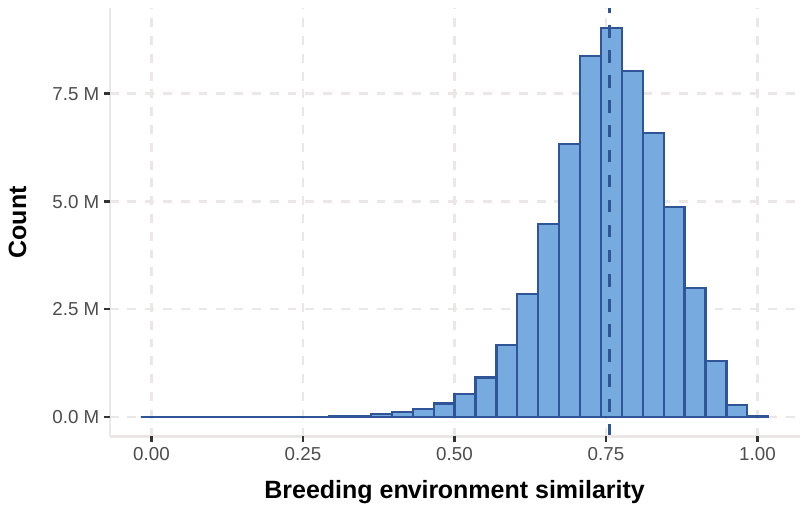

**Supplementary Figure 4 – Breeding environment similarity measure histogram.** Pairwise environmental similarity measure calculated between every pair of birds across all years, using the Euclidean distance of 5 measures of the birds breeding environment (detailed in methods). n = 58,974,720. Median breeding environmental similarity is 0.76.

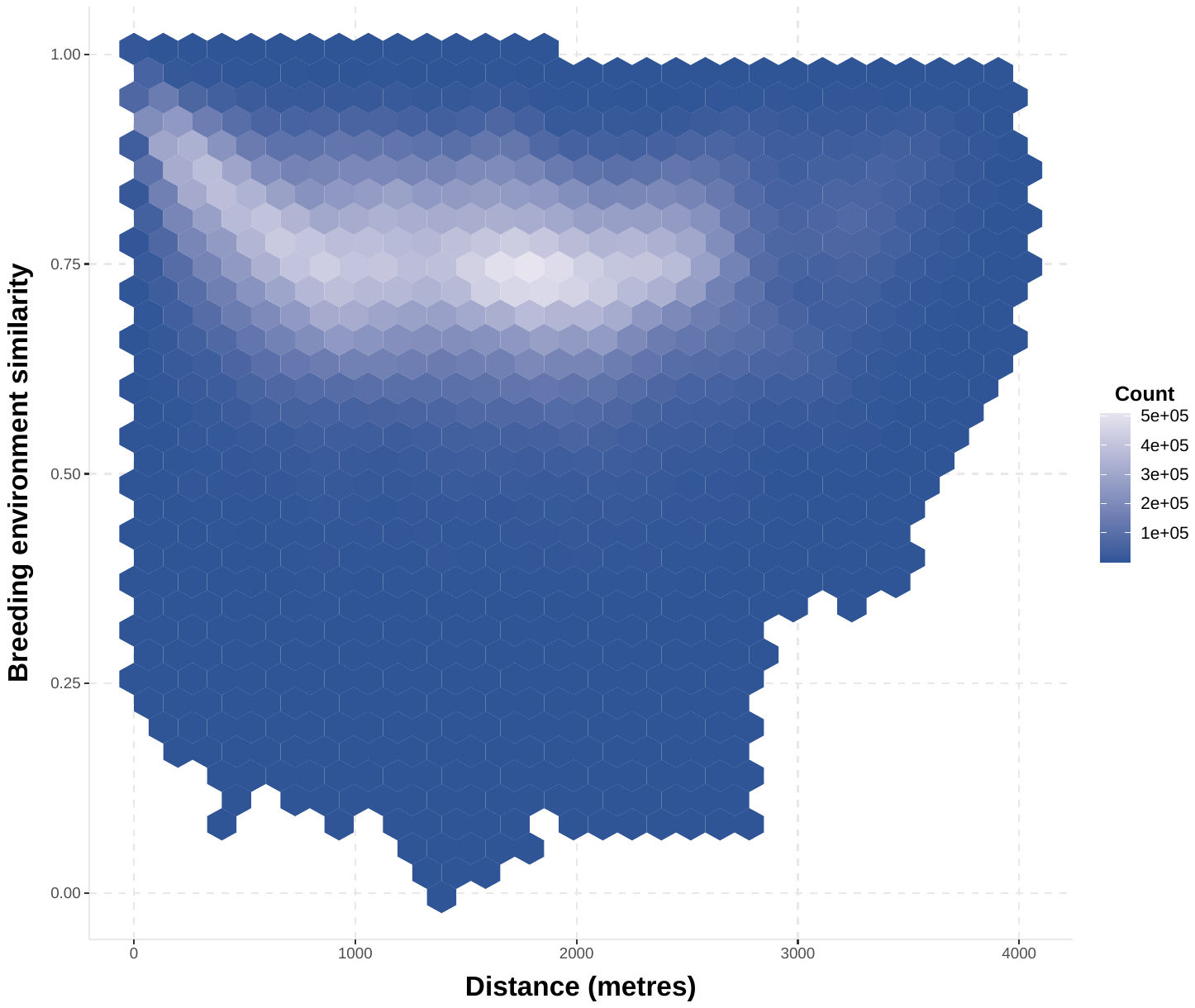

**Supplementary Figure 5 – Breeding environment similarity vs spatial matrix correlation.** Coefficients Relationship between raw distance and breeding environment similarity between all pairwise comparisons of birds (n = 58,974,720). The breeding environment similarity values are calculated by taking a combination of 5 factors of the environment we know to have a relationship with breeding timing, and using Euclidean distance measure in multivariate space between all individuals with every other individual to give one value of similarity for all pairwise comparisons. The final similarity values were again scaled to give a value of 1 along the diagonal, with 0 as the distance between birds in the most dissimilar environments. Mantel tests show a correlation of 0.1923 between the matrix of spatial distance and the matrix of breeding environment similarity.

**Supplementary Table 4 - Fixed effects** Coefficients and standard errors of fixed effects for four models for both traits. Models all include breeding year, additive genetic, permanent environment effects. Nestbox model includes nestbox ID random effect. Spatial models include matrix of spatial proximity random effect and environmental model includes breeding environment similarity matrix random effect.

| *Model* | *Trait* | *Fixed effect* | *Level* | *Coefficient* | *Standard Error* |
| --- | --- | --- | --- | --- | --- |
| **Minimal model** | Laying Date | Female age | Adult | 0.000 | NA |
|  |  |  | Juvenile | 2.386 | 0.100 |
|  | Hatching Date | Female age | Adult | 0.000 | NA |
|  |  |  | Juvenile | 1.540 | 0.023 |
| **Nestbox model** | Laying Date | Female age | Adult | 0.000 | NA |
|  |  |  | Juvenile | 2.387 | 0.099 |
|  | Hatching Date | Female age | Adult | 0.000 | NA |
|  |  |  | Juvenile | 1.536 | 0.023 |
| **Spatial model** | Laying Date | Female age | Adult | 0.000 | NA |
|  |  |  | Juvenile | 2.368 | 0.097 |
|  | Hatching Date | Female age | Adult | 0.000 |  |
|  |  |  | Juvenile | 1.510 | 0.023 |
| **Breed. env. model** | Laying Date | Female age | Adult | 0.000 | NA |
|  |  |  | Juvenile | 2.360 | 0.098 |
|  | Hatching Date | Female age | Adult | 0.000 | NA |
|  |  |  | Juvenile | 1.513 | 0.023 |
